# *tudor-domain containing protein 5-prime* promotes male sexual identity in the *Drosophila* germline and is repressed in females by *Sex lethal*

**DOI:** 10.1101/388850

**Authors:** Shekerah Primus, Caitlin Pozmanter, Kelly Baxter, Mark Van Doren

**Affiliations:** Department of Biology, Johns Hopkins University, Baltimore, Maryland, United States of America

## Abstract

For sexually reproducing organisms, production of male or female gametes depends on specifying the correct sexual identity in the germline. In *D. melanogaster, Sex lethal* (*Sxl*) is the key gene that controls sex determination in both the soma and the germline, but how it does so in the germline is unknown, other than that it is different than in the soma. We conducted an RNA expression profiling experiment to identify direct and indirect germline targets of Sxl specifically in the undifferentiated germline. We find that, in these cells, *Sxl* loss does not lead to a global masculinization observed at the whole-genome level. In contrast, *Sxl* appears to affect a discrete set of genes required in the male germline, such as *Phf7*. We also identify *tudor domain containing protein 5-prime* (*tdrd5p*) as a target for Sxl regulation that is important for male germline identity. *tdrd5p* is repressed by Sxl in female germ cells, but is highly expressed in male germ cells where it promotes proper male fertility and germline differentiation. Additionally, Tdrd5p localizes to cytoplasmic granules with some characteristics of RNA Processing (P-) Bodies, suggesting that it promotes male identity in the germline by regulating post-transcriptional gene expression.

**Author summary:** Like humans, all sexually reproducing organisms require gametes to reproduce. Gametes are made by specialized cells called germ cells, which must have the correct sexual identity information to properly make sperm or eggs. In fruit flies, germ cell sexual identity is controlled by the RNA-binding protein Sxl, which is expressed only in females. To better understand how *Sxl* promotes female identity, we conducted an RNA expression profiling experiment to identify genes whose expression changes in response to the loss of *Sxl* from germ cells. Here, we identify *tudor domain containing protein 5-prime (tdrd5p*), which is expressed 17-fold higher in ovaries lacking *Sxl* compared to control ovaries. Additionally, *tdrd5p* plays an important role in males as male flies that are mutant for this gene cannot make sperm properly and thus are less fertile. Moreover, we find that *tdrd5p* promotes male identity in the germline, as several experiments show that it can shift the germ cell developmental program from female to male. This study tells us that *Sxl* promotes female identity in germ cells by repressing genes, like *tdrd5p*, that promote male identity. Future studies into the function of *tdrd5p* will provide mechanistic insight into how this gene promotes male identity.

## Introduction

Sex determination is an essential process in sexually reproducing species, as the production of eggs and sperm depends on the sexual identity of the germ cells and somatic cells of the gonad. In some animals, such as the medaka fish and the house fly, the sexual identity of the soma determines the sexual identity of the germline. But in other animals, such as fruit flies and mammals, the intrinsic sex chromosome constitution (XX vs. XY) of the germ cells is also important for proper gametogenesis (reviewed in [1]). In such cases, the “sex” of the germ cells must match the “sex” of the soma in order for proper gametogenesis to occur. While studies have revealed a great deal about how sex is determined in the soma, how germline sexual identity is determined by a combination of somatic signals and germline intrinsic properties is much less well understood.

In *Drosophila*, somatic sexual identity is determined by the X chromosome number [2], [reviewed in 3], with two X’s activating expression of the key sex determination gene *Sex lethal* (*Sxl*), promoting female identity. The Sxl RNA-binding protein initiates an alternative RNA splicing cascade to allow female-specific splicing of *transformer (tra*) and, subsequently, *doublesex (dsx*) and *fruitless (fru). dsx* and *fru* encode transcription factors that control somatic sexual identity [reviewed in 4]. *Sxl* is also the key sex determination gene in the germline, as Sxl is expressed in the germline in females, and loss of *Sxl* causes female (XX) germ cells to develop as germline ovarian tumors [5], similar to male (XY) germ cells transplanted into a female soma [6–8]. Further, expression of *Sxl* is sufficient to allow XY germ cells to make eggs when transplanted into a female soma [9]. However, how *Sxl* is activated in the female germline and how it regulates female germline identity remain unknown, except for the fact that both are different than in the soma [6,10–16]. To understand how *Sxl* promotes female germ cell identity, it is essential to discover its targets in the germline.

In this work, we report an RNA expression profiling (RNA-seq) experiment conducted to identify genes regulated downstream of Sxl in the germline. We found that a previously uncharacterized tudor domain containing protein, *tudor domain protein 5-prime (tdrd5p*), is a target of Sxl in the germline. *tdrd5p* is strongly expressed in the *Drosophila* early male germline and is repressed by Sxl activity in the early female germline. It promotes male identity in the germline, and its loss results in germline maintenance and differentiation defects in males, thus reducing their fertility. Tdrd5p protein localizes to cytoplasmic granules related to RNA Processing (P-) Bodies, suggestive of a function in post-transcriptional regulation of gene expression.

## Results

### Analysis of *Sxl*-dependent gene expression changes in the germline

To investigate how Sxl acts to promote female identity in germ cells, we conducted an RNA-seq experiment comparing ovaries with and without *Sxl* function in the germline. To exclude the major gene expression changes that occur during the later stages of gametogenesis, the RNA-seq experiment was done in the *bag of marbles (bam*) mutant background. *bam* is essential for germline differentiation in both males and females; therefore, by using *bam* mutants we focus our experiment on gene expression changes in the early germline, where *Sxl* is expressed most robustly, instead of later stages of gametogenesis (Fig 1A, B). The use of *bam* mutants also gives us the ability to compare similar tissue samples, since ovarian tumors from *bam* mutants and *bam, Sxl* double mutants are very similar [17]. Thus, the two genotypes used for the RNA-seq experiment are both in the *bam*-mutant background, with the experimental genotype knocking down *Sxl* in the germline using RNAi (nanos-Gal4, *UAS-SxlRNAi, bam*), and a control genotype expressing a control RNAi (nanos-Gal4, *UAS-mCherryRNAi, bam*). Sxl proteireads were uniquely mapped to the Drosophila genome, and alln in the germline was dramatically reduced in the *Sxl RNAi* samples relative to controls (Fig 1B, C). Additionally, we conducted RNA-seq on *bam* mutant males compared to females, similar to what has been done previously [18] in order to identify male-enriched vs. female-enriched RNAs in the undifferentiated germline. Libraries were prepared from three biological replicates of each genotype and sequenced with 100bp paired-end reads. The raw reads for all libraries were of very high quality—over 95% of the raw reads received a high quality score. In addition, for each library more than 85% of reads were uniquely mapped to the Drosophila genome, and all replicates had high replicate correlation.

**Fig 1.**
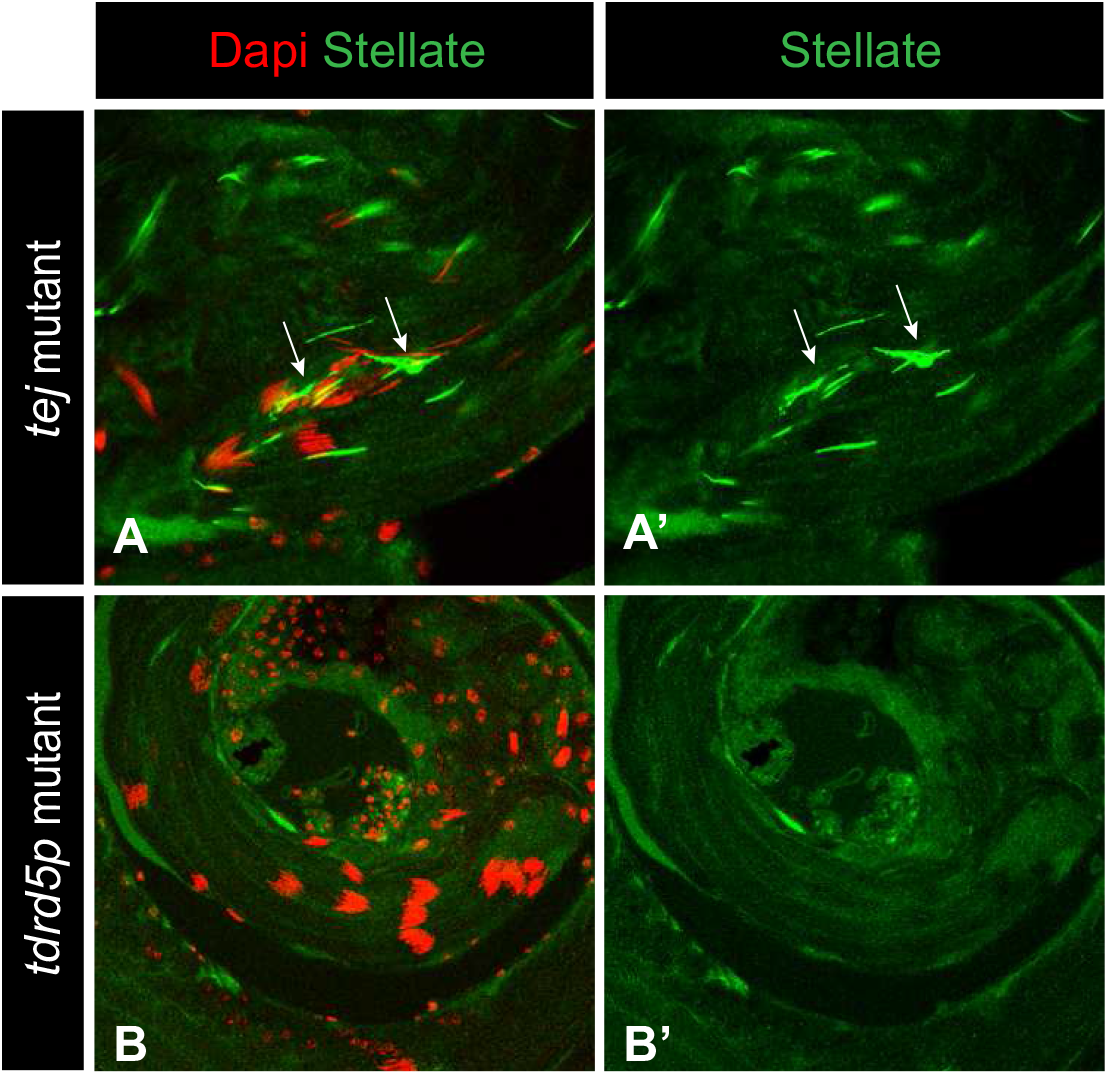
Global effects of *Sxl* loss from the germline. A-C) Immunofluorescence of adult ovaries with antibodies as indicated in the figure. A) wild type ovaries, B-C) ovaries used for the RNASeq experiment: *bam* mutant control RNAi ovaries (B-B’), and *bam* mutant ovaries with *Sxl* knocked down in the germline by RNAi (C-C’). Knockdown of *Sxl* causes a germline tumor phenotype. Note the absence of Sxl from the germline of *Sxl*-RNAi ovaries and presence in all germ cells of the control RNAi. Dashed lines outline ovarioles. Arrows mark the tips of ovarioles. Arrowheads mark differentiating egg chambers. Vasa stains germ cells. D) Principle component analysis comparing each replicate of all four genotypes used in the RNAseq experiment E) Pie chart representing the 94 genes with significant changes in expression levels in *Sxl* RNAi compared to control RNAi and the corresponding changes in *bam* males (bamM) vs. *bam* females.

As Sxl can regulate alternative splicing in the soma, we analyzed our RNA-seq data for changes in exon usage using DEXSeq [19]. We filtered these results for statistical significance (padj < 0.05) and for changes in exon expression that were 2-fold or greater (all DEXSeq data is presented in S1 Table). This identified 44 exons from 34 independent genes. A similar number of exons were upregulated in *Sxl*- (20, 45%) vs. downregulated (24, 55%). However, manual curation of these data did not reveal strong candidates for relevant biological regulation of alternative splicing by Sxl. The main exception was the *Sxl* RNA itself, which in *Sxl RNAi* samples exhibited the characteristic inclusion of the male-specific exon that requires Sxl protein in order to be excluded in the female soma. Thus, the residual *Sxl* RNA present after germline *Sxl* RNAi must be insufficient to produce enough Sxl protein for Sxl autoregulation, further reducing Sxl function in the germline and demonstrating the effectiveness of the germline *Sxl* RNAi approach. This analysis also identified the alternative, testis-specific promoter for the male germline sex determination factor *Plant homeodomain containing protein 7 (Phf7*, [20] as being regulated downstream of *Sxl*, as has previously been described [21]. However, based on this analysis, it does not appear that regulation of alternative splicing is a primary mechanism for Sxl regulation of gene expression in the germline.

We also analyzed changes in overall gene expression levels in the presence of absence of *Sxl* function (DE-Seq [22]). 94 genes were differentially expressed between the two samples (with padj < 0.05), with 40 being upregulated (43%) and 54 being downregulated (57%) in the *Sxl* RNAi samples (S2 Table). 24 of these genes differed 4-fold or more between the two samples (6 upregulated in *Sxl* RNAi, 18 downregulated). We also observed a bias toward X chromosome localization for the differentially expressed genes, with 31.9% being present on the X chromosome compared to only 15.1% of all genes surveyed having X chromosome localization.

We next asked whether Sxl acts as a global regulator of X chromosome gene expression/dosage compensation in the germline, as it does in the soma. Despite the X chromosome bias for differentially expressed genes, relatively few X chromosome genes overall (30 genes or 1.2% of X chromosome loci surveyed) were called as differentially expressed in our analysis. Further, if *Sxl* was a global regulator of X chromosome gene expression, we would expect that the overall level of X chromosome gene expression, relative to autosomal gene expression, would be altered in *Sxl* loss of function. However, the overall ratio of average gene expression between the X chromosome and autosomes was little changed between control and *Sxl* RNAi samples (X:A of 1.19 for controls and 1.24 for *Sxl* RNAi). We conclude that *Sxl* is not a global regulator of X chromosome gene expression in the germline.

Lastly, we evaluated the extent to which loss of *Sxl* in the undifferentiated (*bam-* mutant) female germline leads to masculinization of these cells. We conducted principle component analysis on the individual replicates of all four of our experimental conditions (Fig 1D). We find that *bam Sxl-RNAi* female samples are much more similar to *bam* female samples and control *bam mCherry-RNAi* samples than they are to *bam* male samples. Thus, this analysis does not support a strong masculinization of the *bam-* mutant female germline in the absence of *Sxl*. However, it is also known that *bam* does not affect males and females in exactly the same manner [23] and, in addition, the *bam* male samples also contain male somatic cells while the other samples contain female somatic cells. Both of these factors may contribute to the segregation of the *bam* male sample away from the others. To circumvent this problem, we analyzed the 94 genes differentially expressed between *bam Sxl-RNAi* and control samples to determine whether there was a “male” signature. We determined whether genes differentially expressed in *Sxl-RNAi* compared to controls were also differentially expressed in *bam* males compared to *bam* females. Of the 94 genes differentially expressed in *Sxl* RNAi, a high fraction (44 genes) were also differentially expressed between *bam* male vs. female samples (47%, which is considerably higher than the fraction of total genes in the genome called as different between *bam* male and female, 13%). However, these genes did not always change in the expected direction; only 61% of genes altered in both *Sxl RNAi* and *bam* males changed in the same direction in both, while 39% changed in the opposite direction (Fig 1E). Thus, the *Sxl-RNAi* sample does not appear globally “masculinized” relative to controls and it may be that *Sxl’s* role in repressing male identity in the early germline is restricted to a few specific targets that are important for the male germline.

One such candidate is a previously uncharacterized gene, *CG15930*, that was strongly upregulated in *Sxl-RNAi* ovaries and is normally enriched in testes. *CG15930* exhibits strong homology to mouse Tdrd5 [24], which is essential for male germ cell development and spermatogenesis in mice [25]. However, since *CG15930* is not as similar to Tdrd5 as *Drosophila tejas* [26], and therefore not a paralog of Tdrd5, we named this gene *tudor domain containing protein 5-prime (tdrd5p*). We chose this gene for further study.

### *tdrd5p* is expressed in a sexually dimorphic manner

The RNA-seq expression profiles show that *tdrd5p* has a dynamic expression pattern characteristic of genes with sex-specific functions. *tdrd5p* is 18-fold enriched in *bam* testes compared to *bam* ovaries, and is upregulated 17-fold in *bam*, Sxl-RNAi ovaries relative to *bam*, control-RNAi ovaries (Fig 2A), and is clearly enriched in *bam, Sxl-RNAi* ovaries by RT-PCR (Fig 2B). This is in contrast to *nanos*, which is expressed at similar levels in all of the genotypes in our RNA-seq experiment (Fig 2A), consistent with its role in the germline of both sexes. Changes in *tdrd5p* expression were restricted to total RNA levels, and no change in exon usage was detected. *In situ* hybridization to wild-type gonads revealed that *tdrd5p* expression is highly enriched in the testis, particularly at the apical tip of the testis where the germline stem cells and proliferating gonial cells reside (Fig 2C-D). The finding that *tdrd5p* is expressed at high levels in testes relative to ovaries, and is repressed by *Sxl* in the ovary, suggests that it plays a role in male germline development or function.

**Fig 2.**
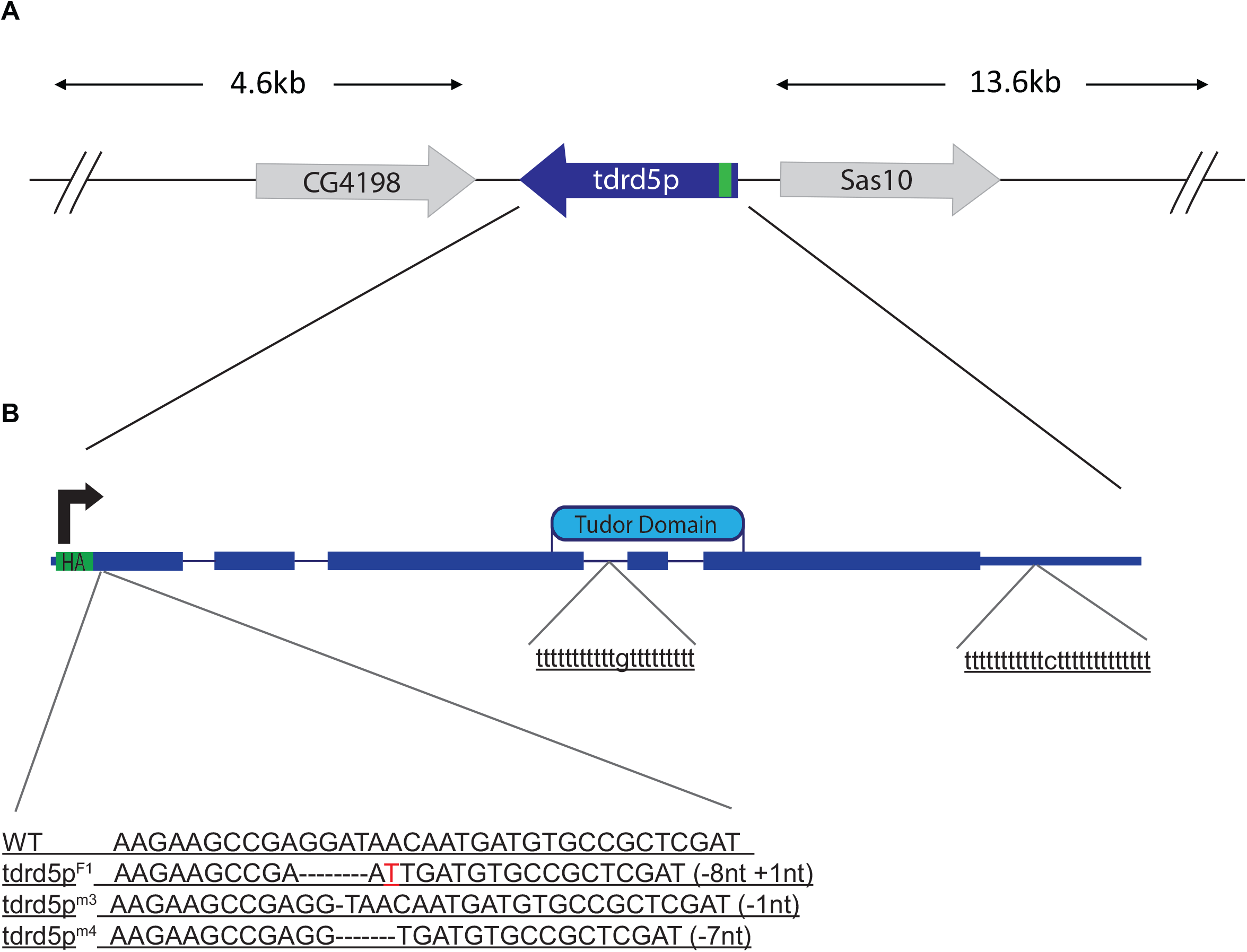
*tdrd5p* has male-biased expression. A) Graph showing the RNA-seq expression profiles of genes with important germline functions. B) RT-PCR showing validation of RNA-seq data for *tdrd5p* upregulation in *Sxl RNAi*. C-D) In-situ hybridization showing expression of *tdrd5p* mRNA in wild type testes (C) and ovaries (D). Sense probe produced no signal (not shown). E-F) Confocal images showing expression of the HA tagged Tdrd5p protein in wild type testes (E) and ovaries (F). Asterisk marks the hub. Arrows show HA:Tdrd5p cytoplasmic punctae. G) Higher magnification image showing size and distribution of HA:Tdrd5p punctae in adult testes. H-H’) Confocal images showing colocalization of HA:Tdrd5p (arrows) with YFP:DCP1 (arrows) in adult testes. Antibodies used are as shown in the figure: HA stains Tdrd5p, YFP stains DCP1, Vasa stains germ cells, TJ stains somatic cells.

To determine the expression pattern of Tdrd5p protein, we generated a genomic transgene that includes a hemagglutinin (HA) epitope tag at the N-terminus of the Tdrd5p protein. The tag was inserted immediately following the start codon of the gene within a 20kb BAC that extends well upstream and downstream of the genes neighboring *tdrd5p* (S1 Fig), and is therefore likely to recapitulate endogenous expression. Anti-HA immunostaining shows that Tdrd5p protein is expressed in the germline of the testis, and is also present in the ovary at lower levels (Fig 2E, F). HA:Tdrd5p is observed in male germline stem cells, spermatogonia and spermatocytes. The protein is seen in distinct foci that are smaller and more numerous in germline stem cells (Fig 2G, arrows), but appear larger in spermatocytes (Fig 2E arrows). The foci are predominantly cytoplasmic, with many abutting a perinuclear germline structure called the nuage. The accumulation of Tdrd5p into cytoplasmic punctae is characteristic of ribonucleoprotein complexes (“RNA bodies”) [reviewed in 27], and suggests that it may be involved in mRNA decay or translational repression. Interestingly, HA:Tdrd5p colocalizes with *decapping protein 1*, (YFP-DCP1, Fig 2H, arrows), which plays a major role in mRNA degradation, and is also required for *osk* mRNA localization to the posterior of the oocyte [28], [reviewed in 29]. As DCP-1 is localized to “Processing bodies” (P-bodies), Tdrd5p appears to be present in a subset of these structures.

HA:Tdrd5p protein expression is upregulated in ovaries that are mutant for *Sxl* function in the germline (Fig 3A-B). Note that the punctae seen in males are also present in *Sxl* RNAi ovaries, though they are fewer in number (Fig 3D, arrows). Thus, the de-repression of *tdrd5p* in *Sxl* RNAi ovaries is also detected at the protein level. Interestingly, the *tdrd5p* mRNA has 2 putative Sxl binding sites [30,31], one within the 3rd intron and the other in the 3’ UTR (S1 Fig). This suggests that Sxl may directly regulate *tdrd5p* expression by binding to one or both of these sites and influencing *tdrd5p* RNA processing in the nucleus or translation in the cytoplasm. To assess Sxl’s direct regulation of *tdrd5p*, we mutated both Sxl binding sites in the a HA:*tdrd5p* transgene (HA:*tdrd5p*^ddel^). We found HA:*tdrd5p*^ddel^ flies show upregulation of HA:Tdrd5p in the female germline (Fig 3C, E), similar to the upregulation caused by loss of *Sxl* from the germline. Quantification of this difference by western blot showed that HA:Tdrd5p is 3-fold upregulated in the female germline of HA:*tdrd5p*^ddel^ flies. However, quantitative RT-PCR analysis specific for the HA-tagged transgenes showed no significant difference in RNA expression between ovaries of HA:*tdrd5p* flies compared to HA:*tdrd5p*^ddel^ flies (data not shown). This suggests that Sxl directly regulates the expression of Tdrd5p protein in the female germline by repressing translation. The basis for the changes we observed in *tdrd5p* RNA levels in *Sxl* RNAi ovaries by RNA-seq remains unknown.

**Fig 3.**
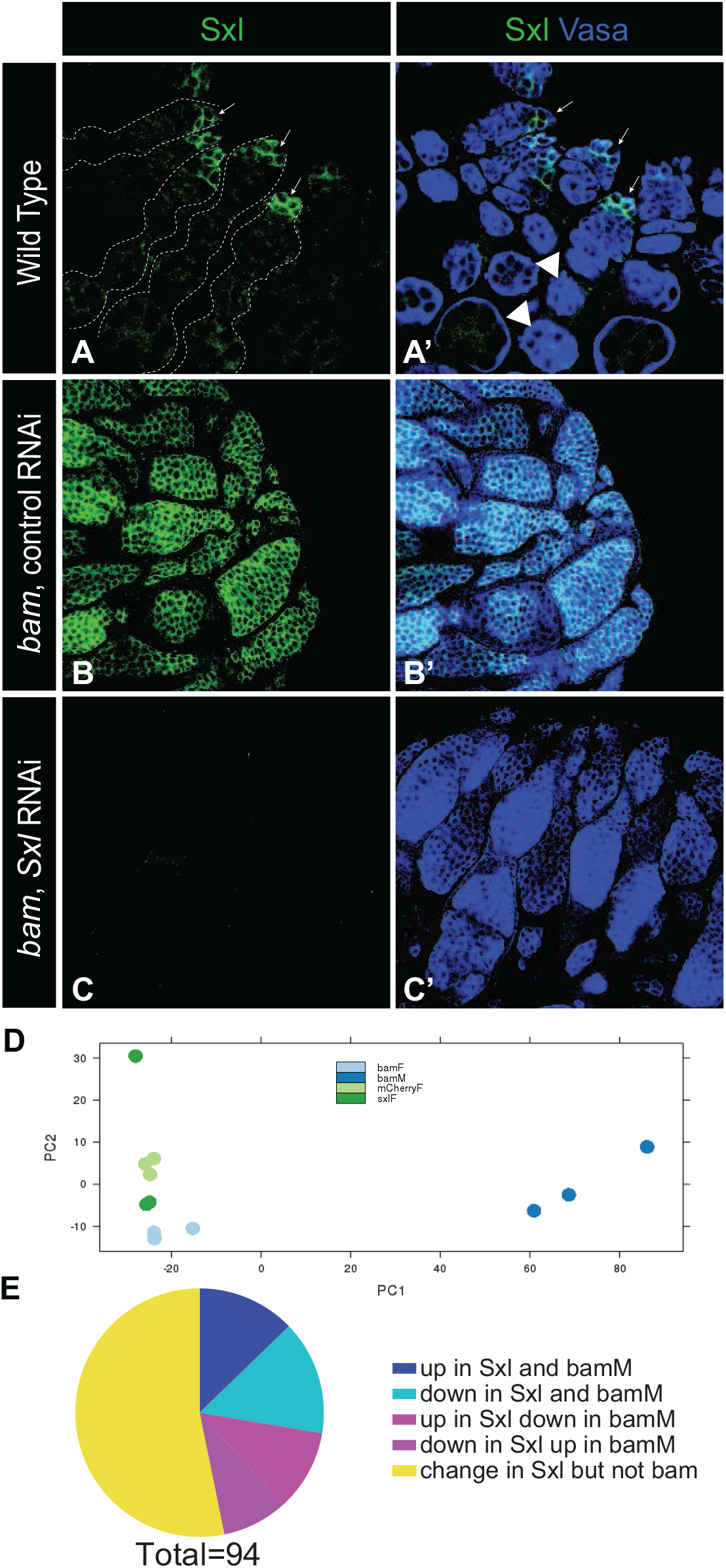
Tdrd5p expression is repressed by *Sxl* function in the female germline. Expression of the HA-tagged Tdrd5p protein in A) wild type ovaries, in B) ovaries with *Sxl* knocked down in the germline, and in C) ovaries expressing the HA:*tdrd5p*^ddel^ transgene, a construct which lacks the two Sxl binding sites. D) Higher magnification image of Sxl-RNAi ovaries showing increase in HA:Tdrd5p punctae (arrows). Antibodies used are as described in figure: HA stains HA:Tdrd5p, Vasa stains germ cells, TJ stains somatic cells. E) Immunoblot comparing HA:Tdrd5p expression in HA:Tdrd5p adult ovaries and HA:Tdrd5p^ddel^ adult ovaries. Antibody against Sxl is used as a loading control.

### *tdrd5p* is required for proper male fertility & germline differentiation

The male-biased expression pattern of *tdrd5p* suggests it may have an important function in the male germline. Knocking down *tdrd5p* function in the germline by RNAi, however, produced no observable phenotype. To conduct a more comprehensive study of *tdrd5p* function we generated *tdrd5p* mutant alleles using CRISPR-Cas9 genome editing. We generated several independent predicted null alleles of *tdrd5p* (S1 Fig), and analyzed male fertility and testis morphology of both young males and aged males. We determined that young (5 days old) *tdrd5p* mutant males have a 50% reduction in fecundity compared to controls (Fig 4F), suggesting that *tdrd5p* is required for proper male fertility.

**Fig 4.**
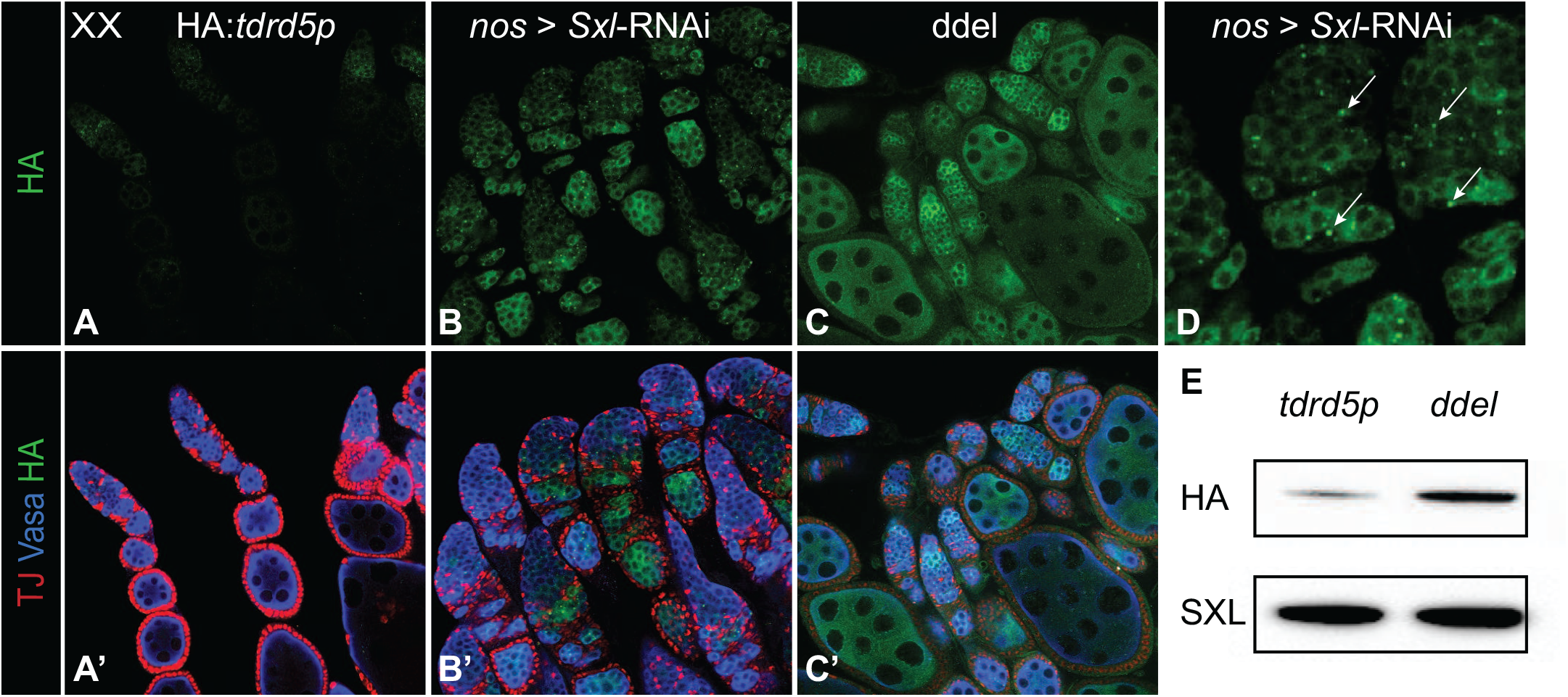
*tdrd5p* is needed for proper male fertility and germline development. A-C”) Confocal images showing germ cell loss phenotype in testes of aged *tdrd5p* mutant flies raised at 25°C (n = 29 testes) and 29°C (n = 39 testes). Asterisk marks the hub. D-E) Confocal images showing hub displacement in *tdrd5p* adult mutant testes. Arrows mark the hub. Vasa stains germ cells, ZFH1 and TJ stain somatic cells, Arm stains membranes, DAPI stains DNA. Antibodies used are as described in the figure. F) Graph showing the total progeny produced per male, comparing *tdrd5p* mutant males to controls. *tdrd5p* mutant males have a 50% reduction in fecundity. Data shown represents the average of two independent experiments with 5 males per test. Significance determined by two-tailed unpaired t test (**P<0.01). See methods for description of control genotype.

To characterize the germ cell defects that may lead to decreased fecundity, we evaluated several aspects of germ cell differentiation in young and aged mutant males. While testes of newly eclosed males appeared similar to wild-type, testes of older males (15-20 days old) exhibited a dystrophic “skinny testis” phenotype with a dramatic reduction in the germline (Fig 4A-C). This was observed in 7% of animals raised at 25°C and 18% of animals raised at 29°C. Additionally, 15% of males exhibited a displaced hub phenotype (Fig 4E arrow), where the hub is no longer located at the apical tip of the testis as seen in wild-type (Fig 4D arrow). These combined phenotypes suggest a defect in proper germline differentiation and maintenance. Analyzing the expression of critical germ cell differentiation genes such as *bam* and *spermatocyte arrest* (required for meiotic cell cycle progression) [32] did not shed further light on the germ cell defect. Therefore, while the morphological defects present at a low penetrance, their overall effects culminate into a substantial reduction in fecundity; a phenotype which supports *tdrd5p* importance in male germline development.

Tudor domain-containing proteins have well known functions in small RNA pathways, transcriptional regulation, and the assembly of snRNPs [reviewed in 33]. The closest *Drosophila* homolog to *tdrd5p* is *tejas*, and the closest mammalian homolog to *tdrd5p* is mouse TDRD5. Both *tejas* and TDRD5 have been shown to function in the piRNA pathway and to repress the expression of transposons in the germline [25,26,34]. Interestingly, we found no changes in transposon expression in *tdrd5p* mutants in our RNA-seq analysis (data not shown). We also analyzed the expression of a wide variety of transposons by quantitative rtPCR and found little to no difference between wt and tdrd5p-mutant testes (data not shown). Defects in the piRNA pathway can also lead to increased accumulation of the Stellate protein in the male germline [35,36]. We examined the expression of Stellate in *tdrd5p* mutant males, as well as in *tdrd5p* mutant males also heterozygous for mutant alleles of *tejas, ago3* or *aubergine* (two PIWI proteins with key functions in the piRNA pathway). None of these testes showed the increased Stellate expression observed in *tejas* homozygous mutant testes (S2 Fig, [26]). Therefore, while *tdrd5p’s* activity is important for the proper development of the male germline, it has a distinct function from regulation of transposon expression.

### *tdrd5p* promotes male identity in the germline

The sex-specific nature of *tdrd5p’s* expression and mutant phenotype suggests it may play a role in promoting male germline sexual identity. However, unlike the male-specific *Phf7* gene [20], expression of UAS-*tdrd5p* in the female germline did not, by itself, result in defects in the female germline (data not shown). To further investigate *tdrd5p’s* role in sexual identity, we decided to conduct our experiments using the sensitized genetic backgrounds frequently used for the investigation of genes involved in sexual identity. Females mutant for *transformer* (*tra*)—a key player in the somatic sex determination pathway—undergo a transformation so that the somatic gonad of XX *tra* mutants develops as male instead of female. However, because the germline is XX, and therefore incompatible with spermatogenesis, the germline of these testes is highly undeveloped, causing these animals to be sterile (Fig 5B). A strong test of a gene’s ability to promote male identity in the germline is to determine whether it is sufficient to induce XX germ cells to enter spermatogenesis in these animals. Indeed, expression of *tdrd5p* in the germline of XX *tra* mutants resulted in a robust rescue of spermatogenesis. 18% of these animals had highly developed testes that were wild type in size, containing all of the stages of germ cell differentiation up to spermatocytes (Fig 5C, note: since these animals lack a Y chromosome and the spermatogenesis genes located there, they were not expected to be fertile). This is strong evidence signifying that *tdrd5p* promotes male identity in the germline.

**Fig 5.**
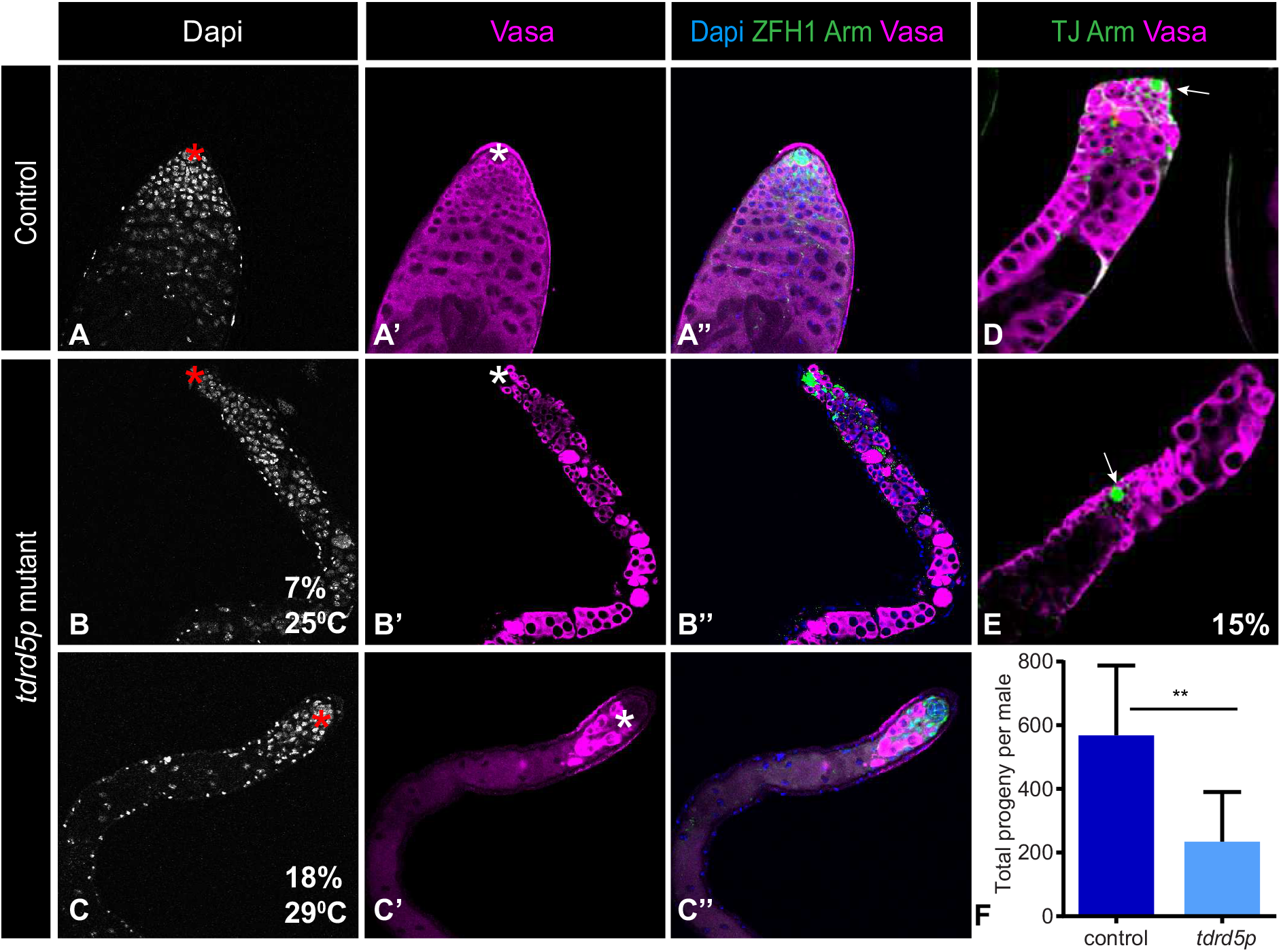
*tdrd5p* promotes male identity in the germline. Confocal images of A) wild type XY adult testes, B) XX *tra* mutant adult testes, and C) XX *tra* mutant adult testes rescued by *tdrd5p* expression in the germline, driven by *nos*-gal4-VP16 driver (n=101 testes). D-F’) Ectopic expression of *tdrd5p* enhances the *snf* mutant phenotype. Confocal images of D) *snf*/+ adult ovaries, E) *snf*/+ adult ovaries with ectopic expression of *tdrd5p*, driven in the germline by *nos*-gal4-VP16, and F) *snf/snf* mutant adult ovaries, showing the germline tumor phenotype. Vasa stains germ cells, TJ stains somatic cells, Arm and HTS stain membranes, Dapi stains DNA, antibodies used are as indicated in the figure. G) Quantification of the degree of enhancement of the *snf* and *otu* mutant phenotypes with ectopic *tdrd5p* expression. The number of ovaries scored per genotype is shown in each bar.

Similarly, if *tdrd5p* promotes male identity in the germline, we would expect that it would be able to enhance the ability of other mutations to masculinize the female germline. Homozygous mutants of *ovarian tumor (otu*) and *sans fille (snf*) have been shown to masculinize the female germline and cause germline tumors in ovaries, similar to *Sxl*-RNAi [37–40], while females heterozygous for mutant alleles of *otu* and *snf* are fully fertile and have normal ovary morphology. However, ectopic expression of *tdrd5p* in the germline of females heterozygous for *otu* or *snf* resulted in the formation of ovarian tumors. 25% of *snf/+; nos* > *tdrd5p* ovaries have large, pervasive germline tumors similar to the homozygous *snf* mutants (Fig 5D-F). Another 25% of these ovaries show a complete loss of the germline (Fig 5G), which phenocopies strong sex determination mutants [37,41]. Additionally, 40% of *otu*/+; *nos* > *tdrd5p* ovaries exhibit either ovarian tumors or complete loss of germline. This evidence supports *tdrd5p* as a male-promoting factor in the germline. We therefore conclude that *tdrd5p* functions in germline sexual identity; it promotes male identity in the germline and is repressed by *Sxl* in the female germline.

## Discussion

### The role of *Sxl* in the germline

It has been known for many years that *Sxl* is necessary for female germline identity [7,8], and *Sxl* has also been shown to be sufficient to allow XY germ cells to undergo oogenesis [9]. It is likely that *Sxl* plays multiple roles in the germline, both to promote female identity in the early germline, perhaps as early as in the embryonic germline [9], and in regulating the differentiation of the germline during oogenesis [42]. Here we examined the role of *Sxl* more specifically in the undifferentiated germline through the use of *bam* mutants. Principle component analysis indicated that, under these conditions, samples with *Sxl* function reduced in the germline clustered close to control female samples, and far from male samples. While some of the male/female differences may be contributed by the somatic cells present in these samples, we conclude that reducing *Sxl* function in the undifferentiated germline does not lead to a dramatic masculinization at the whole-genome level. In contrast, we propose that the role for *Sxl* in the early germline may be restricted to a relatively small number of changes in sex-specific germline gene expression that are important for female vs. male germline function.

Recently, a genomic analysis of ovaries mutant for the RNA splicing factor *sans fille (snf*) was conducted [21]. This is considered to also be a *Sxl* germline loss-of-function condition as one important change in *snf*-mutant ovaries is a loss of *Sxl* expression and an ovarian tumor phenotype that can be rescued by *Sxl* expression [17,42,43]. In contrast to our results, an increased expression of spermatogenesis genes was observed in *snf* tumorous ovaries compared to wild type ovaries. It is likely that changes in these “differentiation” genes were not observed in our *bam*-mutant samples since germline differentiation is arrested at an earlier stage in *bam* mutants, allowing us to focus on the undifferentiated germline. Thus, these two analyses can help separate the role of *Sxl* in regulating early germline sexual identity vs. later aspects of sex-specific germline differentiation. Interestingly, one “differentiation” gene was identified in both RNA-seq analyses: the testis-specific basal transcription factor *TATA Protein Associated Factor 12L (Taf12L* or *rye*). This may indicate that *Taf12L* could play a role in the undifferentiated germline as well as the later stages of spermatocyte differentiation. In addition, both analyses found evidence for differential regulation of the important male germline identity factor *Phf7* [20], where an upstream promoter is utilized preferentially in the male germline and is repressed downstream of *Sxl* in females [21 and data not shown]. This indicates a role for *Phf7* in both the early and differentiating germline. Finally, we did not observe strong candidates for targets of alternative RNA splicing regulated by Sxl. The only strong candidate for alternative RNA splicing was the *Sxl* RNA itself, where the male-specific exon was retained in the residual *Sxl* RNA from the *Sxl RNAi* samples. This provides further evidence that *Sxl* autoregulation occurs in the germline as it does in the soma, as has previously been proposed [44]. It is likely that Sxl may also act at the level of translational control in the germline, as our evidence indicates here for regulation of Tdrd5p. Future experiments to identify Sxl-associated germline RNAs will be important for investigating this mechanism of action, as has recently been conducted [9].

In addition to its role in sex determination in the soma, *Sxl* also acts to initiate global X chromosome gene regulation and dosage compensation through translational control of *male-specific lethal-2* [45], and it is possible that *Sxl* plays a similar role in the germline. Whether or not the germline even undergoes dosage compensation is controversial, and thoughtful work has led to opposite conclusions [46,47]. Further, if dosage compensation does exist in the germline, it must utilize a separate mechanism from the soma, as the somatic dosage compensation complex members *msl1* and *msl2* are not required in the germline [48,49]. However, *Sxl* could retain an *msl*-independent role to regulate global X chromosome gene expression in the germline. We did not observe evidence for this. First, few X chromosome genes were differentially expressed between *Sxl-* and control samples (30 genes or 1.2% of X chromosome genes tested). Second, the ratio of average gene expression between X chromsome genes and autosomes was very similar in *Sxl-* females compared to control females (X/A for controls: 1.24, *Sxl*-: 1.19) and this was similar when considering only genes in particular expression categories (e.g. all genes with some expression in both samples). Thus, it appears likely that *Sxl’s* role in the germline is distinct from that in the soma; it acts to control sex-specific gene regulation and sexual identity in both the germline and the soma, but acts as a general regulator of X chromosome gene expression and dosage compensation only in the soma.

### *tdrd5p* promotes male germline identity

We show here that *tdrd5p* is both a target for *Sxl* regulation and is important for male germline identity and spermatogenesis. *tdrd5p* expression is highly male-biased, both at the RNA and protein levels (Fig 2). When female germ cells are sensitized by partial loss of female sex determination genes, expression of *tdrd5p* exacerbates the masculinized phenotype in these germ cells (Fig 5). Significantly, expression of *tdrd5p* is sufficient to promote spermatogenesis in XX germ cells present in a male soma (XX *tra-* mutant testes, Fig 5). Thus, *tdrd5p* clearly has a role in promoting male germline identity.

Loss of *tdrd5p* also has a strong effect on male fecundity, even though it is not absolutely required for spermatogenesis. The 50% reduction in fecundity is a strong effect and indicates that *tdrd5p* plays an important role in the male germline. However, the fact that some spermatogenesis still proceeds suggests that other factors act in combination with *tdrd5p* to control this process. One good candidate is *Phf7* which we have previously demonstrated to have a similarly important, but not absolute, requirement for spermatogenesis [20]. However, our analyses of *tdrd5p, Phf7* double mutants did not reveal any synergistic effect on male germline development or fecundity (data not shown). This suggests that other important players in promoting male germline identity remain to be identified.

Our data indicate that Tdrd5p regulates male germline identity by influencing post-transcriptional gene regulation. Other tudor-domain containing proteins have been shown to act in RNA-protein bodies to influence RNA stability and translational regulation [reviewed in 33]. Further, Tdrd5p localizes to cytoplasmic punctae, specifically a subset of the punctae that also contain Decapping Protein1 (DCP-1), suggesting that these bodies are related to Processing bodes (P-bodies) that are known to control post-transcriptional gene regulation [28,50], [reviewed in 29]. Interestingly, Tdrd5p’s closest homologs, Tejas in flies and TDRD5 in mice, have been shown to regulate piRNA production and transposon regulation [25,26]. Further, their localization to VASA-containing nuage is thought to influence transposon control [26,51]. However, we observe no changes in transposon expression regulated by *tdrd5p*, and Tdrd5p does not co-localize with VASA in nuage. We have also not observed any genetic interaction between *tejas* and *tdrd5p* (data not shown). Thus, we propose that Tdrd5p plays a distinct role in regulating male germline identity and spermatogenesis, and this may be in the regulation of mRNAs rather than transposons. One possibility is that the role of mouse TDRD5 in transposon regulation and spermatogenesis has been separated into the roles of Tejas in transposon regulation and Tdrd5p in regulating male identity and spermatogenesis in flies. It is widely known that regulation of germline identity is dependent on post-transcriptional mechanisms involving tudor-domain proteins such as the original Tudor protein [52], which helps define the germ plasm, an RNA body that regulates germline identity [reviewed in 53]. Further, the regulation of sex-specific gametogenesis is also dependent on RNA bodies and their requisite tudor-domain proteins, such as TDRD5 in mouse [54], [reviewed in 27]. Our studies indicate that initial germline sexual identity is similarly regulated by post-transcriptional mechanisms, including RNA bodies containing other, distinct, TUDOR-domain proteins such as Tdrd5p.

## Materials and methods

### Library Generation and sequencing

Gonads were dissected from 1-3 day old flies raised at 25°C. Ovaries were dissected from virgin females. 3 biological replicates were dissected for each genotype. Total RNA was isolated from all genotypes using RNA-bee (Tel-Test). Contaminating DNA was removed from the RNA using Turbo-DNA-free (Ambion). 200ng of RNA was used to prepare each library using the illumina TruSeq RNA Library Prep kit v2. 100bp paired-end read sequencing was done by the Johns Hopkins Genetic Resources Core Facility. The *bam* mutant male and female libraries were sequenced in one lane and the *Sxl*-RNAi, control-RNAi libraries were sequenced in separate lane, therefore having 6 libraries per lane.

### Read mapping, quality, and differential expression analysis

Quality of raw reads was assessed using the fastQC kit (Babraham Institute). RNA-Seq reads were mapped to the Drosophila genome using Ensembl BDGP6 release 85, and Bowtie 2.2.9, TopHat 2.1.1, and HTSeq 0.9.1 [55–57]. Differential gene expression analysis was done using DESeq using ensemble annotation BDGP6 [22]. Adjusted P value of 0.05 used for significance cutoff. Differential exon analysis was done using DEXSeq [19].

### Fly stocks and Fecundity Tests

The fly stocks used were obtained from Bloomington Stock Center unless otherwise indicated. *bam*^1^ [58], *bam^Δ86^* (BDSC# 5427), *nos*-Gal4 [59], the control RNAi used was p{VALIUM20-mCherry}attP2 (BDSC# 35785), uas-*Sxl*-RNAi=TRiP.HMS00609 (BDSC# 34393), uas-*CG15930*-RNAi=TRiP. GL01046 (BDSC# 36882), *Snf^148^, otu^17^, tej^48-5^*, attP40{*nos*-Cas9} (NIG-FLY# CAS-0001), YFP:dDCP1 (a kind gift from Ming-Der Lin, [28]). Fecundity tests were carried out by setting up crosses with one *tdrd5p* mutant male and 15 virgin females of the control stock. The control stock used is *nos*-Cas9 isogenized to FM7KrGFP fly stock to reproduce the treatment of the *tdrd5p* mutant fly lines while screening for transformants. Each male was mated with virgin females for 4 days. Females were then discarded and each male was placed with another 15 virgin females in a new bottle. This was repeated twice more for a total of four mating bottles per male. All offspring were counted by day 18. Total offspring per male was calculated by averaging the number of offspring from each of the four mating bottles for each male.

### Immunofluorescence

Adult ovaries and testes were fixed, blocked and stained as previously described [60]. All images were taken with a Zeiss LSM 510 confocal microscope.

Primary antibodies and the concentrations used are as follows: chicken anti-Vasa 1:10,000 (K. Howard); rabbit anti-Vasa 1:10,000 (R. Lehmann); mouse anti-Sxl 1:8 (M18, DSHB); mouse anti-Armadillo 1:100 (N2 7A1, DSHB); rat anti-HA 1:100 (3F10, Roche); guinea pig α-TJ (1:1,000; generated by J. Jemc using the same epitope as previously described [61]); mouse anti-HTS 1:4 (1B1, DSHB). DSHB: Developmental Studies Hybridoma Bank. Secondary antibodies were used at 1:500 (Alexa-fluor). Samples were mounted in vectashield mounting solution with DAPI (vector Industries).

### RT-PCR, quantitative RT-PCR & In-situ hybridization

For RT-PCR and qRT-PCR, total RNA was isolated from ovaries and testes using RNA-bee (Tel-Test). Contaminating DNA was removed from the RNA using Turbo-DNA-free (Ambion). RNA was converted to cDNA using Superscript II (Invitrogen). qRT-PCR was done using 2 biological replicates and in technical triplicate, using SYBR green detection. In-situ hybridization was carried out as previously described [62]. DIG-labelled sense and antisense probes were synthesized by in vitro transcription of PCR product generated from RP98-1M22 BAC (BACPAC Resources Center).

### Mutagenesis and BAC-tagging

Mutant alleles of *tdrd5p* were created using CRISPR-Cas9 mediated genome editing. Small guide RNA (sgRNA) was designed and cloned following the Perrimon lab protocol [63], using the U6b-sgRNA-short vector described therein. The sgRNA was injected by Best Gene inc into *nos*-Cas9(II-attP40) flies. The HA:*Tdrd5p* transgenic flies were generated by BAC recombineering [64,65], using the CH322-188C18 BAC obtained from the BACPAC Resources Center. A 3xHA epitope tag was added to the N-terminus of the gene (S1 Fig). This construct was also modified to delete the Sxl binding sites in the intron and the 3’UTR. The Sxl binding site in *tdrd5p* 3’UTR was deleted using QuickChange II site-directed mutagenesis kit (agilent). The intronic binding site was removed by deleting the entire intron.

## Acknowledgements

We thank the fly community for providing stocks and reagents, either directly as specified in the materials and methods or through their contributions to the Bloomington Stock Center, the Developmental Studies Hybridoma Bank, and BACPAC Resources. We would also like to thank David Mohr and the Johns Hopkins Sequencing facility for technical expertise on high throughput RNA sequencing, Dr. Frederick Tan and other members of the Johns Hopkins Community for technical expertise on Bioinformatics and sequencing analysis, as well as members of the Chen, Johnston and Van Doren laboratories for helpful discussions. Imaging was performed at the Integrated Imaging Center at the John Hopkins University.

## Supporting Information

**S1 Table. RNAseq analysis_changes in exon usage**.

**S2 Table. RNAseq analysis_gene expression changes**.

**S1 Fig.**
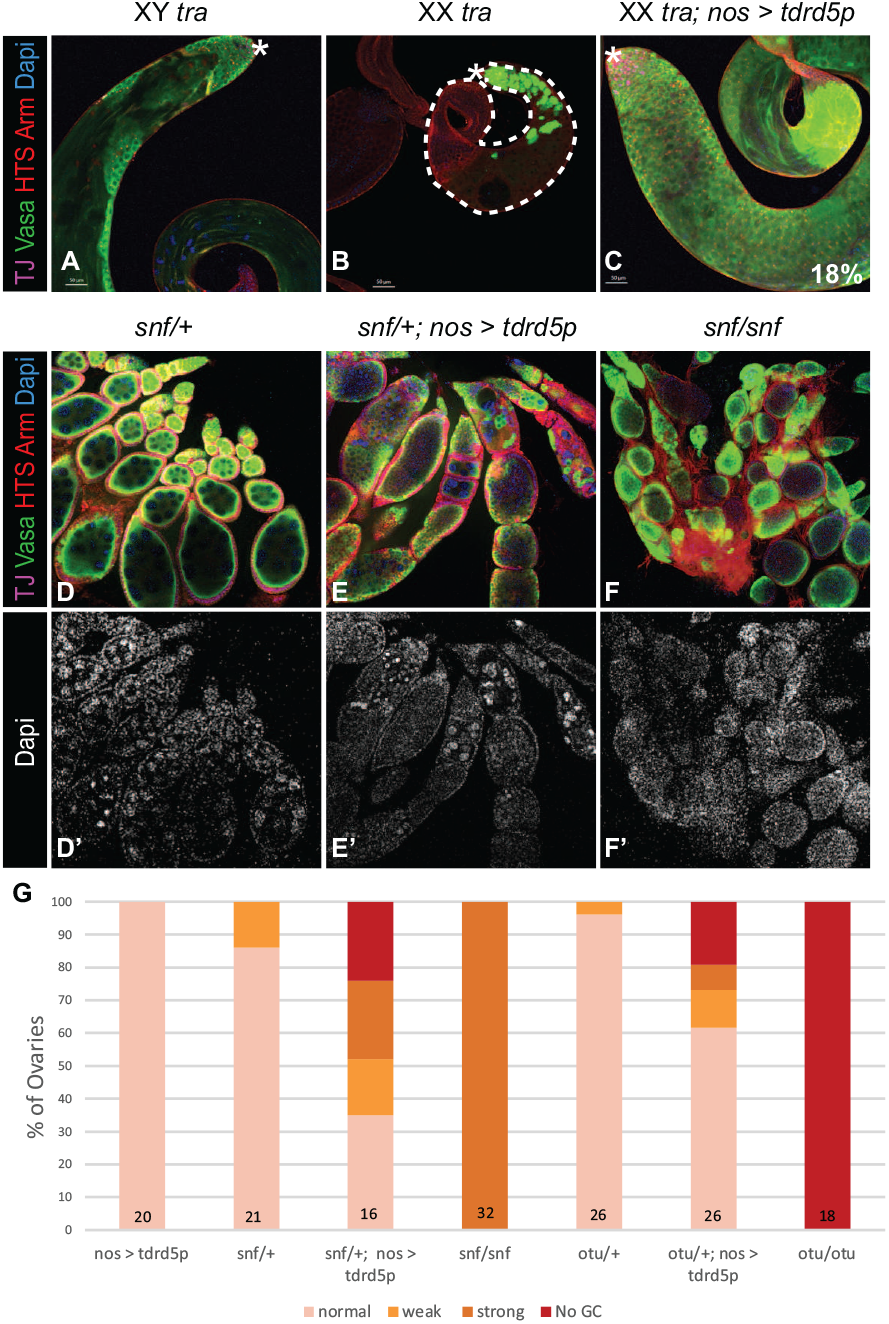
*tdrd5p* gene model. *A*) Schematic of the Bacterial Artificial Chromosome used to insert a hemagglutinin (HA) tag in *tdrd5p* by BAC recombineering (not drawn to scale). The *tdrd5p* gene is shown in reference to its closest neighboring genes: sas10 at its 5’ end and CG4198 at its 3’ end, which are both on the opposite strand. B) Gene model of *tdrd5p*. The HA tag was inserted at the N-terminus of *tdrd5p* immediately after the start codon. Putative Sxl binding sites are located in the 3^rd^ intron and the 3’UTR. Mutant alleles described were generated by CRISPR-Cas9-mediated gene editing in the 1^st^ exon.

**S2 Fig.**
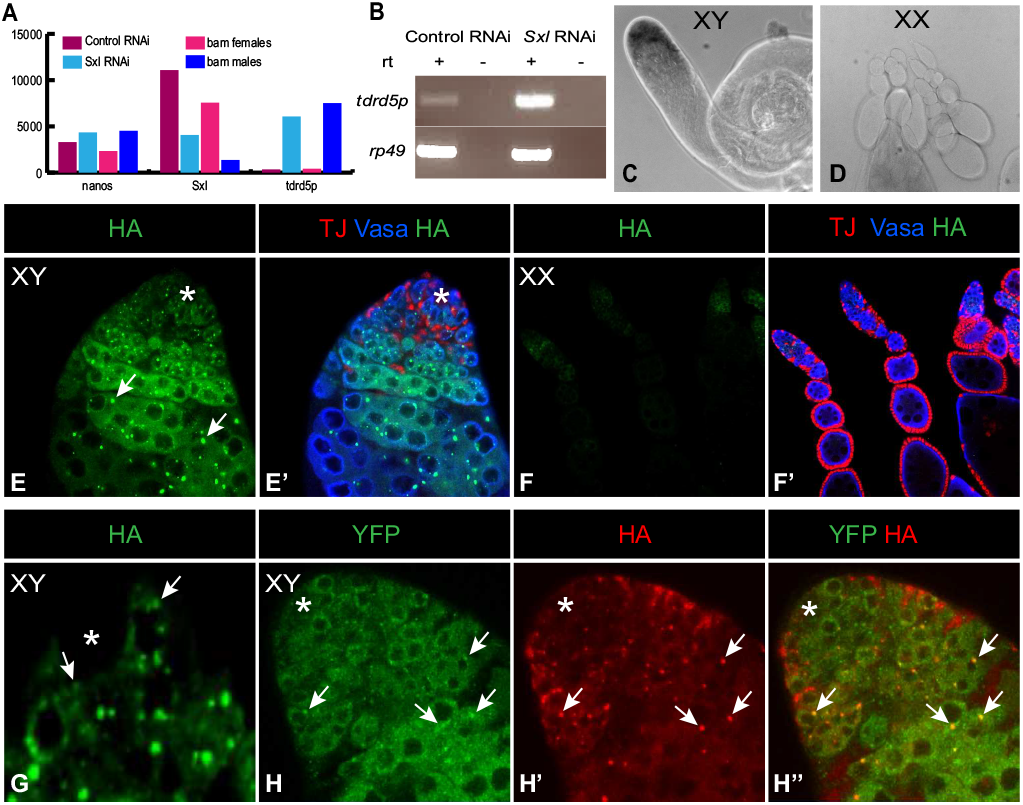
Stellate transposon is not upregulated in *tdrd5p* mutants. Confocal images showing expression of stellate crystals (arrows) in A) *tej* mutant testes but not in B) *tdrd5p* mutant testes. Antibodies used are as indicated in the figure.

